# The assembly of Hfq into foci-like structures in response to long-term nitrogen starvation in *Escherichia coli*

**DOI:** 10.1101/2020.01.10.901611

**Authors:** Josh McQuail, Amy Switzer, Lynn Burchell, Sivaramesh Wigneshweraraj

## Abstract

The initial adaptive responses to nutrient depletion in bacteria often occur at the level of RNA metabolism. Hfq is an RNA-binding protein present in diverse bacterial lineages and contributes to many different aspects of RNA metabolism. We demonstrate that Hfq forms a distinct and reversible focus-like structure in *E. coli* specifically experiencing long-term nitrogen (N) starvation. Using the ability of T7 phage to replicate in N starved bacteria as a biological probe of *E. coli* cell function during N starvation, we demonstrate that Hfq foci have a role in the adaptive response to long-term N starvation. We further show that Hfq foci formation does not depend on gene expression during N starvation and occurs independently of the N regulatory protein C (NtrC) activated initial adaptive response to N starvation. The results serve as a paradigm to demonstrate that bacterial adaptation to long-term nutrient starvation can be spatiotemporally coordinated and can occur independently of *de novo* gene expression during starvation.

**Significance Statement:** Bacteria have evolved complex strategies to cope with conditions of nitrogen (N) adversity. We now reveal a role for a widely studied RNA binding protein, Hfq, in the processes involved in how *Escherichia coli* copes with N starvation. We demonstrate that Hfq forms a distinct and reversible focus-like structure in long-term N starved *E. coli*. We provide evidence to suggest that the Hfq foci are important features required for adjusting *E. coli* cell function during N starvation for optimal adaptation to long-term N starvation. The results have broad implications for our understanding of bacterial adaptive processes in response to stress.

## Introduction

Bacteria in their natural environments seldom encounter conditions that support continuous growth. Hence, many bacteria spend the majority of their time in a state of little or no growth, because they are starved of essential nutrients including carbon, nitrogen and transitional metals. To maximise chances of survival during prolonged periods of nutrient starvation and facilitate optimal growth resumption when nutrients become replenished, bacteria have evolved complex adaptive strategies. Bacteria initially respond to nutrient deficiency by remodelling gene expression through the synthesis and degradation of RNA. Nitrogen (N) is an essential element of most macromolecules in a bacterial cell, including proteins, nucleic acids and cell wall components. Thus, unsurprisingly, when *Escherichia coli* cells experience N starvation, they attenuate growth and elicit nitrogen stress response (Ntr response). The Ntr response is activated by the transcription factor N regulatory protein C (NtrC) concomitantly with N run out. By directly activating the transcription of *relA*, the gene responsible for the synthesis of the major bacterial stress signalling nucleotide guanosine pentaphosphate, NtrC couples the Ntr response with the stringent response (1) and thereby induces a rapid and large-scale reprogramming of cellular processes to adjust to N starvation. The Ntr response primarily involves the synthesis of proteins associated with transport, and assimilation of nitrogenous compounds into glutamine and glutamate, either catabolically or by reducing the requirement for them in other cellular processes (1–4).

Small regulatory (non-coding) RNA molecules (sRNAs) play an important part regulating the flow of genetic information in response to nutrient starvation in many bacteria (5–9). sRNAs can base pair with target mRNAs leading to enhanced translation or inhibition of translation and/or alteration of mRNA stability (10, 11). In order to form productive interactions with target mRNAs, most sRNAs require an RNA binding protein (RBP). In many bacteria of diverse lineages, the RBP Hfq plays a central and integral role in sRNA mediated control of gene expression. Emerging results now reveal that Hfq has diverse functions in bacteria that expands beyond its widely understood role in catalysing sRNA-mRNA base pairing. Hfq is also involved in ribosomal RNA processing and assembly of functional ribosomes (12); tRNA maturation (13); and regulation of RNA degradation (14–17). Recently, Hfq was shown to contribute to the distribution of sRNAs to the poles of *E. coli* cells experiencing envelope stress, suggesting a role for Hfq in spatiotemporal regulation of gene expression (18). Though Hfq is widely studied in the context of bacterial stress responses, its role in the Ntr response is unknown. Hence, the initial aim of this study was to investigate the role of Hfq in the post-transcriptional regulation during the Ntr response. However, unexpectedly, we uncovered a property of Hfq, which appears to occur independently of the Ntr response, but one that is important for adapting to long-term N starvation.

## Results and Discussion

### Absence of Hfq compromises the ability of *E. coli* to survive N starvation

To determine if Hfq has a role in the adaptive response of *E. coli* to N starvation, we grew a batch culture of wild-type (WT) and Δ*hfq E. coli* in a highly defined minimal growth media with a limiting amount (3 mM) of ammonium chloride (NH_4_Cl) as the sole N source (19). Under these conditions, when NH_4_Cl (i.e. N) in the growth medium runs out (N-; Fig. 1*A*), the bacteria enter a state of complete N starvation, and subsequent growth attenuation (19). As shown in Fig. 1*A*, the initial growth rate (μ) of WT (μ=64.0±0.30 min/generation) and Δ*hfq* (μ=67.3±1.28 min/generation) *E. coli* under N replete conditions (N+; Fig. 1*A*) did not differ greatly. However, as the ammonium chloride levels became depleted (ND; Fig. 1*A*), the growth rate of the Δ*hfq* (μ=71.3±3.45 min/generation) dropped by ~8% relative to WT bacteria (μ=66.1±0.26 min/generation). Both strains attenuated growth at N-. We then measured the number of colony forming units (CFU) in the population of WT and Δ*hfq* bacteria as a function of time under N starvation. As shown in Fig. 1*B*, the proportion of viable cells in the WT population moderately increased over the initial 24 h under N starvation but gradually declined as N starvation ensued beyond 24 h. This initial moderate increase in the portion of viable cells over the initial 24 h was not observed for Δ*hfq* bacteria (Fig. 1*B*). In contrast, the portion of viable cells in the Δ*hfq* population rapidly decreased as N starvation ensued. For example, after 24 h of N starvation (N-24), only ~27% of the Δ*hfq* mutant population was viable compared to the WT population (Fig. 1*B*). After 168 h under N starvation (N-168), the majority (~99%) of bacteria in the Δ*hfq* population were nonviable relative to bacteria in the WT population (Fig. 1*B*). The viability defect of Δ*hfq* bacteria was partially reversible to that of WT levels when *hfq* was exogenously supplied via a plasmid (Fig. 1*B*, *inset*).

**Fig. 1.**
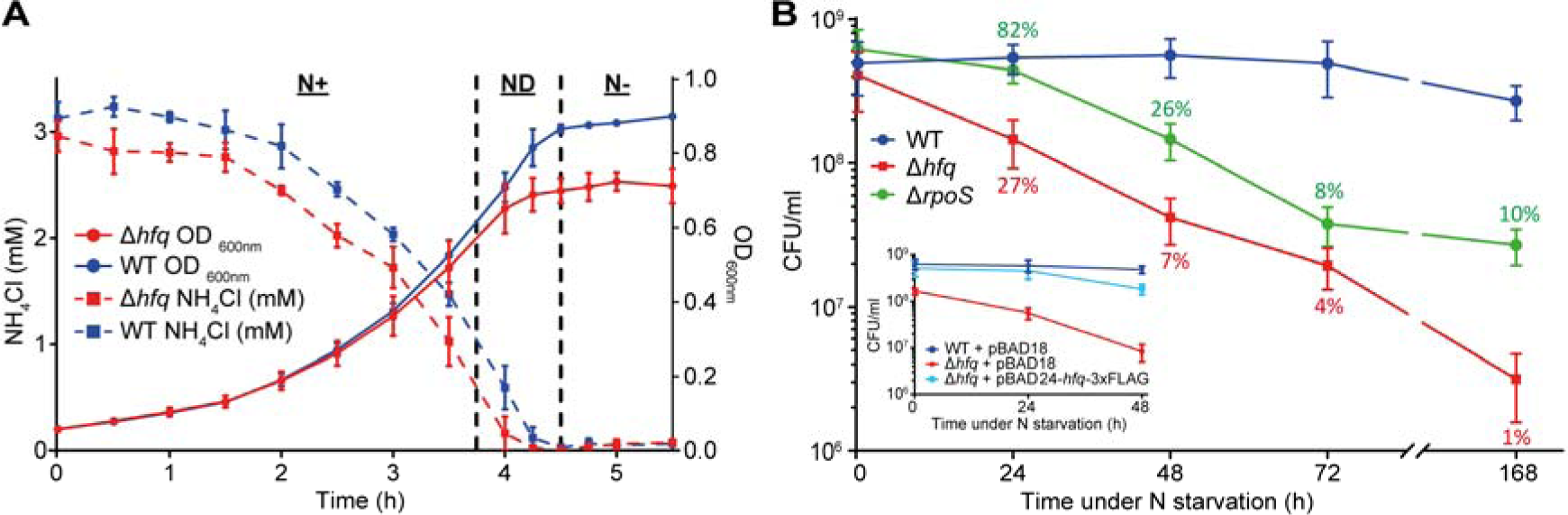
Absence of Hfq compromises the ability of *E. coli* to survive N starvation. (*A*) Growth and NH_4_Cl consumption of WT and Δ*hfq E. coli* grown in N limited conditions. Error bars represent standard deviation (n=3). (*B*) Viability of WT, Δ*hfq* and Δ*rpoS E. coli* during long-term N starvation, measured by counting CFU. Inset shows viability of WT and Δ*hfq E. coli* complemented with plasmid-borne *hfq* (pBAD24-*hfq*-3xFLAG) measured by counting CFU. The percentage of Δ*hfq* and Δ*rpoS* viable cells compared that of WT cells are indicated. Error bars represent standard deviation (n□= □3).

As Hfq is a positive regulator of expression of *rpoS* (20–23), the RNAP promoter-specificity factor (σ^S^), which is responsible for the transcription of diverse stress response associated genes, we considered whether the inability of Δ*hfq* bacteria to adjust their metabolism to cope with N starvation is due to compromised σ^S^ activity. To investigate this, we calculated, as above, the number of CFU in the population of Δ*rpoS* bacteria as a function of time under N starvation. The results revealed that, following 24-48 h of N starvation, the Δ*rpoS* bacteria were significantly better at surviving N starvation than Δ*hfq* bacteria (Fig. 1*B*). For example, at N-24, ~82% of Δ*rpoS* were viable relative to WT bacteria. In contrast, at N-24, only ~27% of the Δ*hfq* bacteria were viable. After 48 h under N starvation, the Δ*rpoS* bacteria displayed a better ability to survive N starvation than the Δ*hfq* bacteria (Fig. 1*B*). Overall, the results demonstrate that the absence of Hfq compromises the ability of *E. coli* to survive N starvation in a manner that, at least partly, independent of Hfq’s role in the regulation of *rpoS*.

### Absence of Hfq compromises the ability of T7 to replicate in long-term N starved *E. coli*

As the T7 phage can infect and replicate in exponentially growing and stationary phase (i.e. nutrient starved) *E. coli* cells equally well (24), we used T7 as a biological probe of *E. coli* cell function during N starvation to evaluate the metabolic capacity and capability of the N starved cellular environment of WT and Δ*hfq* bacteria to support T7 replication. Put simply, we reasoned that as T7 heavily relies on bacterial resources for replication, any perturbations to these resources due to a dysfunctional ability to adjust to N starvation (i.e. in Δ*hfq* bacteria) could have a negative impact on the efficacy of T7 replication. We thus compared the time it took for T7 to decrease the starting density (OD_600nm_) of the culture of WT bacteria at the onset of N starvation (N-) and at N-24 by ~50% (T_lysis_) following infection. We did this by resuspending bacteria from N- and N-24 in media containing ~3 mM NH_4_Cl and T7 phage (NH_4_Cl was added to re-activate cellular processes that might be required for T7 infection, but might have become repressed upon N starvation). As shown in Fig. 2*A*, the T_lysis_ of a culture of WT bacteria at N- and N-24 was ~62 min and ~106 min, respectively. The T_lysis_ of Δ*hfq* bacteria infected at N-was delayed by ~17 min compared to WT bacteria and this resulted in moderate growth of the Δ*hfq* bacterial culture before cell lysis was detectable (Fig.2*A*). Strikingly, however, T7 replication was substantially compromised in Δ*hfq* bacteria infected at N-24 (Fig. 2*A*) and detectable lysis of Δ*hfq* bacteria at N-24 was delayed by ~83min compared to WT bacteria. This delay in lysis of Δ*hfq* bacteria at N-24 was partially reversible to that seen in WT bacteria when *hfq* was exogenously supplied via a plasmid (Fig. 2*B*).

**Fig. 2.**
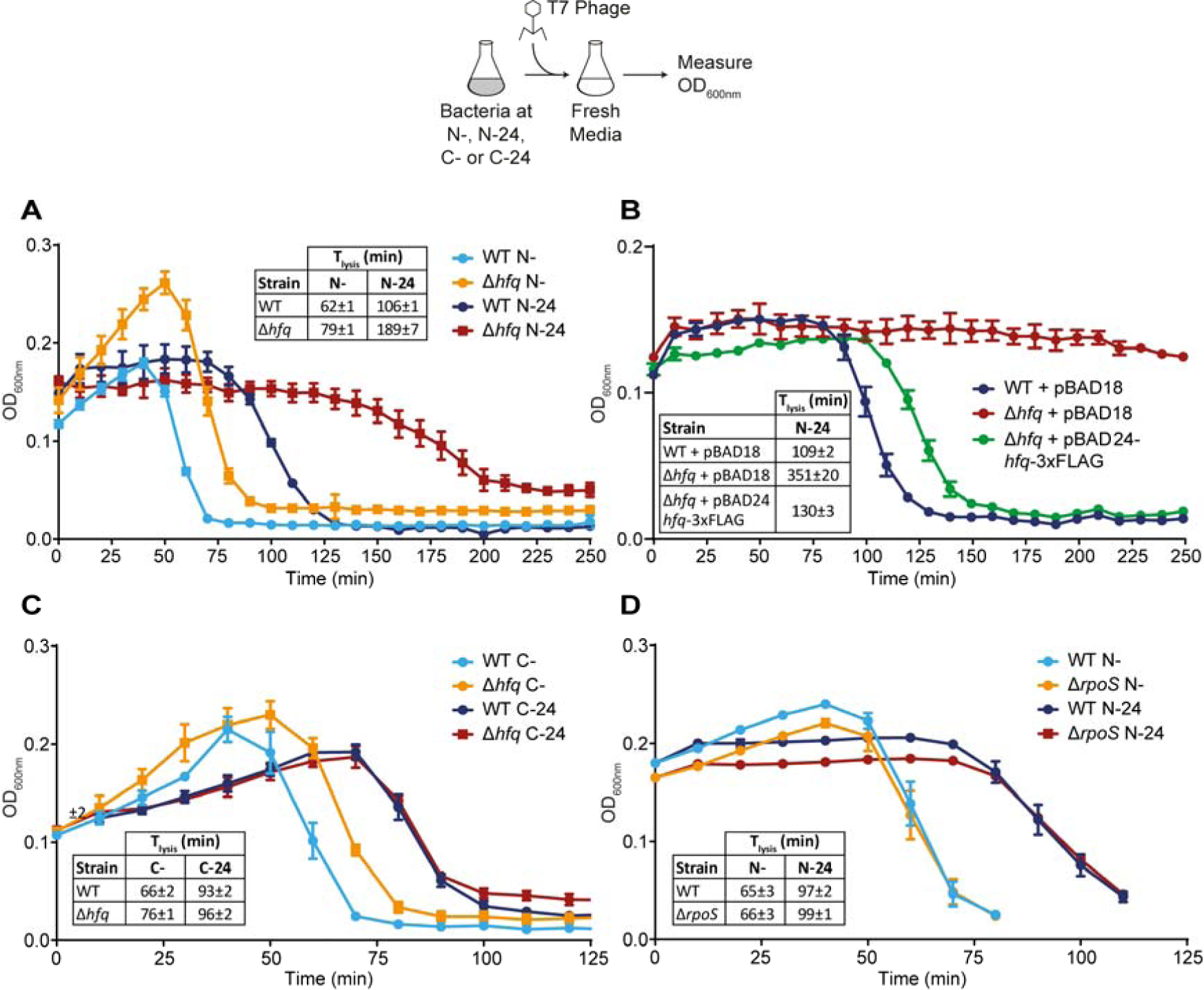
Absence of Hfq compromises the ability of T7 to replicate in long-term N starved *E. coli*. (*A*) Graph showing the optical density (OD_600nm_) as a function of time of WT and Δ*hfq E. coli* cells from N- and N-24 following infection with T7 phage. The time taken for the starting OD_600nm_ value of the culture to decrease by ~50% (T_lysis_) is indicated in the inset table. (*B*) As in (*A*), but the Δ*hfq E. coli* cells were complemented with plasmid-borne *hfq* (pBAD24-*hfq*-3xFLAG). (*C*) As in (*A*), but experiments were conducted with C starved bacteria. (*D*) As in (*A*), but experiments were conducted with Δ*rpoS E. coli* cells. Error bars represent standard deviation (n□= □3).

We next investigated whether the observed difference in T_lysis_ was specific to N starvation. Hence, we measured T_lysis_ in carbon (C) starved WT and Δ*hfq* bacteria. This was achieved by growing bacteria for 24 h in minimal growth media with limiting amount of C (0.06% (w/v) of glucose instead of 0.4% (w/v) of glucose) and excess NH_4_Cl (10 mM instead of 3mM). Under these conditions, the bacteria run out of C and thus become C starved before the source of N is completely consumed. As shown in Fig. 2*C*, the dynamics of lysis of the bacterial culture immediately following onset of C starvation (C-) and N-were indistinguishable. However, interestingly, experiments with 24 h C starved bacteria (C-24), did not produce a difference in T_lysis_ between WT and Δ*hfq* bacteria (Fig. 2*C*) as seen with N-24 bacteria (Fig. 2*A*). Clearly, the compromised ability of T7 to replicate in N-24 bacteria is specific to N starvation. The results also suggest that the compromised ability of T7 to replicate in the Δ*hfq* strain was not due to a requirement of T7 for Hfq for replication *per se* and that the ability of T7 to replicate in N starved *E. coli* is a faithful reporter of *E. coli* cell function during N starvation. Additional experiments with the Δ*rpoS* strain revealed that the compromised ability of T7 to replicate in Δ*hfq* bacteria at N-24 was not due to an indirect effect of the absence of Hfq on *rpoS* (Fig. 2*D*). This is consistent with the view that the loss of viability of Δ*hfq* bacteria as a function of time under N starvation was not due to loss of *rpoS* activity (Fig. 1*B*). Overall, we conclude that the compromised ability of T7 to replicate in long-term N starved Δ*hfq* bacteria is due to a dysfunctional ability of mutant bacteria to adapt to long-term N starvation.

### Hfq molecules assemble into foci-like structures in long-term N starved *E. coli* cells

To better understand the role of Hfq during long-term N starvation in *E. coli*, we used photoactivated localisation microscopy combined with single-molecule tracking to study the intracellular behaviour of individual Hfq molecules in live *E. coli* cells. To do this, we constructed an *E. coli* strain containing photoactivatable mCherry (PA-mCherry) fused C-terminally to Hfq at its normal chromosomal location. Control experiments established that the ability of (i) PA-mCherry-tagged Hfq bacteria to survive N starvation was indistinguishable from that of WT bacteria (Fig. S1) and (ii) T7 phage to replicate in 24 h N starved bacteria containing PA-mCherry-tagged Hfq was indistinguishable from that of WT bacteria (Fig. S2). These results thus indicate that the presence of the PA-mCherry tag on Hfq did not compromise its function under our experimental conditions. We used the apparent diffusion coefficient (*D*\*) of individual Hfq-PAmCherry molecules, calculated from the mean squared displacement of trajectories, as a metric for the single molecule behaviour of Hfq. As shown in Fig. 3*A* and 3*B*, in bacteria at N+ and N-, the *D*\* values of Hfq molecules were largely similar (though we note that our *D\** values were ~6-fold lower than those measured by Persson et al (25) with Dendra2-tagged Hfq in exponentially growing *E. coli* in lysogeny broth, our measurements are largely consistent with those measured by Park et al (26) with mMaple3 tagged Hfq in *E. coli* growing in defined in minimal media). In bacteria at N-24, we detected a large increase in the proportion of molecules with a lower *D*\*. Strikingly, this was due to Hfq forming a *single* focus-like feature (~360 nm in diameter), which was present usually, but not exclusively, at a cell pole (Fig. 3*C*). These features, hereafter referred to as the Hfq foci, were seen in ~90% of the cells from N-24 that we analysed. The *D*\* value of the majority (~75%) Hfq molecules *within* the Hfq foci was <0.08 (Fig. S3). We collated the *D*\* values of multiple fields of view (typically containing ~50-300 bacterial cells per field of view) to maximise our data pool. We then used a *D*\* of <0.08 as a threshold to define the relatively immobile population of Hfq molecules within the bacterial cells within the field of view imaged (typically ~50-300 bacterial cells). We then calculated the proportion of all Hfq molecules within this immobile population as a percentage of total number of tracked Hfq molecules within all the bacterial cells within the same field of view imaged to derive a value (%H_IM_) to indirectly quantify the efficiency of foci formation by Hfq under different conditions. In other words, cells containing detectable Hfq foci will have an increased %H_IM_ compared to cells without detectable Hfq foci. According to this criteria, the %H_IM_ values were ~14, ~16, and ~44 in bacteria at N+, N- and N-24, respectively.

**Fig. 3.**
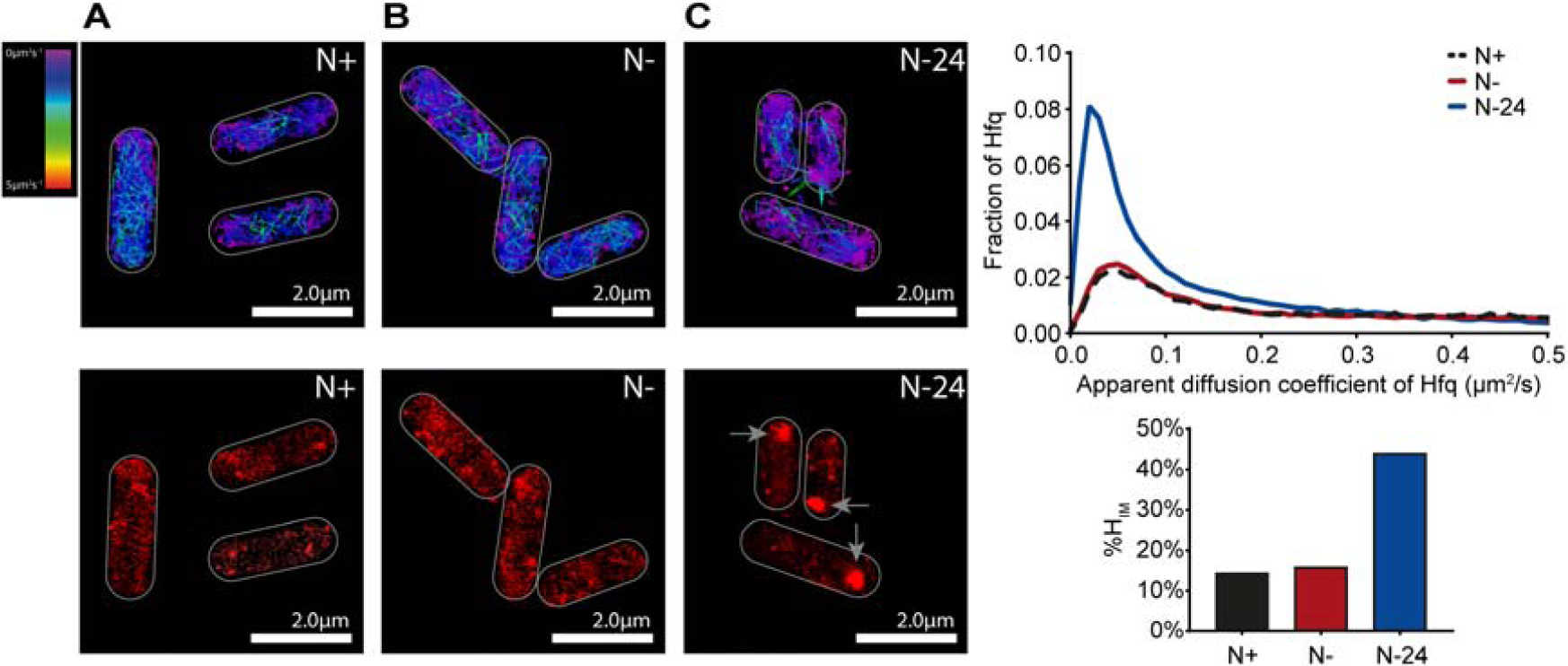
Hfq molecules assemble into focus-like structures in long-term N starved *E. coli* cells. Representative single molecule tracks (*top panel*) and PALM images (*bottom panel*) of Hfq in *E. coli* cells from N+, N- and N-24. In the PALM images, the arrows indicate the Hfq foci. The graph shows the distribution of the apparent diffusion coefficient of Hfq molecules and the bar chart shows %H_IM_ values (see text for details).

The Hfq foci, once formed, persisted for at least 168 h under N starvation (Fig. S4). We did not detect any foci under identical experimental conditions in N-24 bacteria in *E. coli* strains with PAmCherry fused to RNA polymerase, MetJ (the DNA binding transcriptional repressor of genes associated with methionine biosynthesis, which is similar in size to Hfq), or ProQ (a new class of sRNA binding protein in bacteria) (27–29) (Fig. S5). This result suggested that foci formation is a biological property specific to Hfq. Further, we did not detect Hfq foci in bacteria that were starved for C for 24 h (Fig. S6). Similarly, we did not detect Hfq foci in 24 h old stationary phase cultures grown in standard lysogeny broth (Fig. S7). These results suggest that Hfq formation is a phenomenon specific to N starvation. Consistent with this view, we also observed Hfq foci formation in 24 h old cultures grown in media containing 3 mM L-glutamine, D-serine or L-aspartic acid as the sole N source (Fig. S8). These results indicate that Hfq foci formation is a response to depletion of N source and not a response restricted to the depletion of NH_4_Cl. As shown in Fig. S9, the intracellular levels of Hfq did not increase over 24 h under N starvation (when foci have formed). This suggests that the Hfq foci seen in *E. coli* from N-24 are unlikely to be due to accumulation of Hfq as N starvation ensued. To determine whether the foci represent aberrant aggregates of Hfq molecules, we obtained the fraction of aggregated proteins in bacteria from N+, N- and N-24 (as described in (30)) and attempted to identify Hfq by immunoblotting with anti-PAmCherry-tag antibodies following separation of the samples on a sodium dodecyl sulphate polyacrylamide gel. As shown in Fig. S9, we did not detect Hfq in any of the fractions containing aggregated proteins (i.e. in the insoluble fraction). This suggests that the Hfq foci are unlikely to be aberrant aggregates of Hfq molecules. Finally, Fortas et al (31) previously showed that the unstructured C-terminal region of Hfq has the intrinsic property to self-assemble, albeit into long amyloid-like fibrillar structures, *in vitro*. Further, related studies by Taghbalout et al (32) and Fortas et al (31) showed that Hfq forms irregular clusters in exponentially growing *E. coli* cells in lysogeny broth and that the C-terminal region of Hfq was required for this clustering behaviour of Hfq, respectively. To investigate whether the C-terminal amino acid residues of Hfq contribute to foci formation, we constructed an *E. coli* strain with a PAmCherry tag fused to Hfq at its normal chromosomal location, but which had amino acid residues 73-102 deleted (Hfq_Δ73-102_). As shown in Fig. S10, foci formation by WT Hfq and Hfq_Δ73-102_ did not markedly differ in bacteria at N-24. We note that a study by Vecerek et al showed that a larger truncation of the C-terminal region (Hfq_Δ65-102_) resulted in a mutant protein that was defective in some but not all of its functions (33, 34). However, consistent with the result that foci formation by Hfq_Δ73-102_ and WT Hfq is indistinguishable, the Hfq_Δ73-102_ variant survived N starvation as well as WT bacteria (Fig. S10). This suggests that the Hfq foci are different to Hfq clusters/aggregates previously seen *in vitro* and *in vivo*. Overall, we conclude that the majority of Hfq molecules assemble into a single focus-like structure in *E. coli* cells experiencing long-term N starvation.

### Hfq foci formation contributes to adapting *E. coli* cell function as N starvation ensues

As the results thus far indicate that Hfq foci formation happens in long-term N starved *E. coli*, we next investigated the dynamics of Hfq foci formation during the first 24 h of N starvation. At the same time, we also assessed the ability of T7 to replicate in WT and Δ*hfq* bacteria during the first 24 h of N starvation (by measuring the difference in T_lysis_ (ΔT_lysis_) between WT and Δ*hfq* bacteria as shown in Fig. 2) to determine whether the time point during long-term N starvation when detectable Hfq foci form in WT bacteria coincides with the compromised ability of T7 to replicate in Δ*hfq* bacteria (where Hfq foci cannot form). As shown in Fig. 4*A*, no Hfq foci were detected in bacteria that had been N starved for up to 3 hours (i.e. at N-3). However, in bacteria that have been starved of N for ~6 h (N-6), we began to detect clustering of Hfq resembling Hfq foci and by N-12 discernible Hfq foci were clearly seen (Fig. 4*A*). Strikingly, a substantial increase in ΔT_lysis_ occurred concomitantly with the assembly of Hfq into the foci (Fig. 4*A* and Fig. 4*B* compare N-9 with N-12, 18 and 24). We conclude that that Hfq foci formation is a process that occurs gradually over the course of N starvation in *E. coli* and that Hfq foci contributes to adapting *E. coli* cell function as N starvation ensues.

**Fig. 4.**
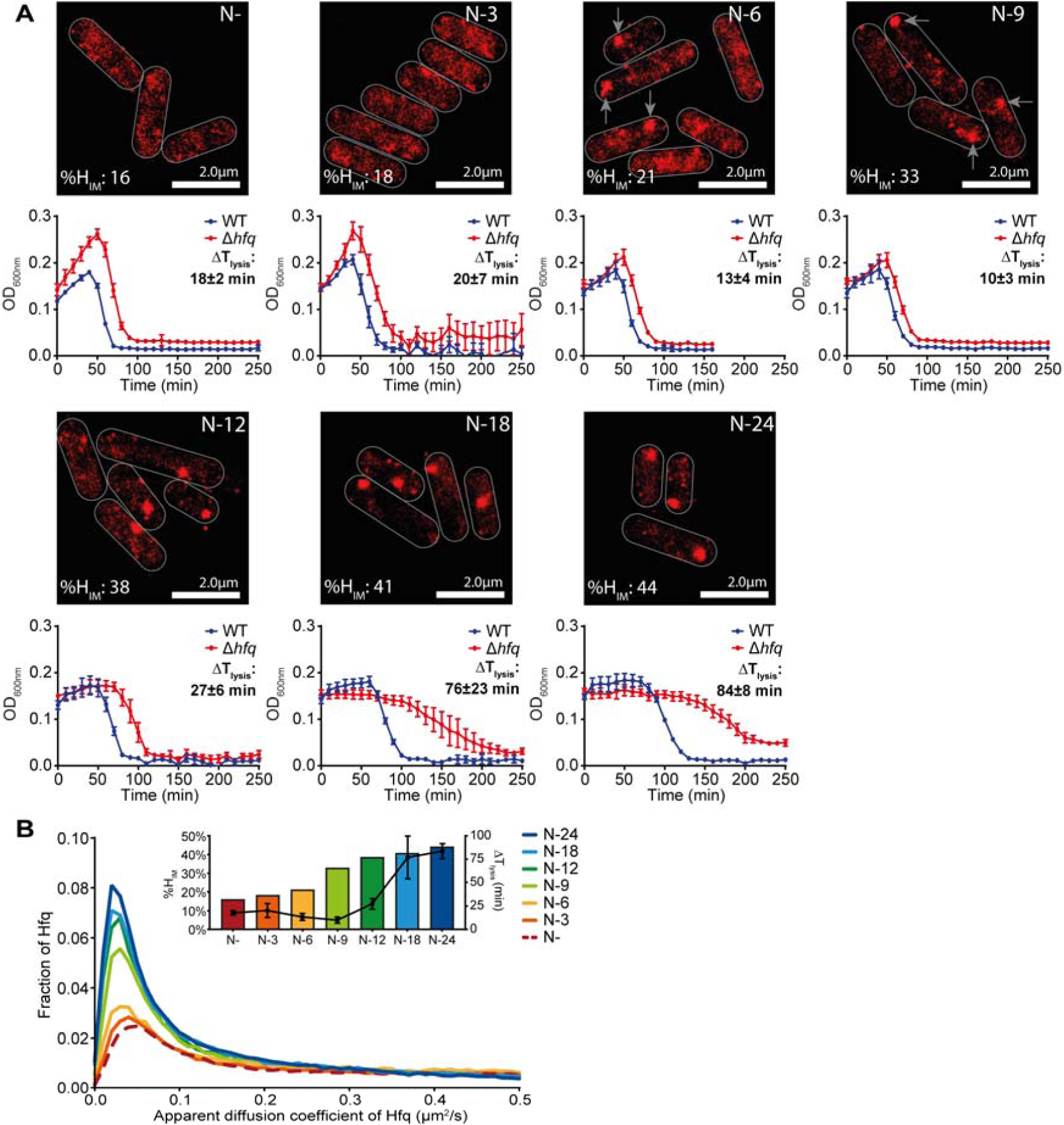
Hfq foci formation contributes to adapting *E. coli* cell function as N starvation ensues (*A*) Representative PALM images of Hfq in *E. coli* cells as a function of time under N starvation (*top panel*). Images were taken at the indicated time points and the arrows point to the Hfq foci. Graphs (*bottom panel*) with the optical density (OD_600nm_) as a function of time of WT and Δ*hfq E. coli* cells following infection with T7 phage are shown below each corresponding PALM image. The difference in time taken for the OD_600nm_ of the culture to decrease by ~50% (T_lysis_) between WT and Δ*hfq* is indicated (ΔT_lysis_). (*B*) Graphs showing the distribution of apparent diffusion coefficient of Hfq molecules at the different sampling time points and the corresponding %H_IM_ and ΔT_lysis_ values are shown as the inset (see text for details).

### Hfq foci are reversible

We next considered that if Hfq foci formation is a direct response to long-term N starvation, then the foci should dissipate when long-term N starved bacteria are replenished with N. To explore this, we harvested N starved (hence growth attenuated) bacteria at N-24 and inoculated them into fresh growth media. This, as expected, resulted in the resumption of growth and we detected the dissipation of the Hfq foci just ~1 h after inoculation in fresh N replete growth medium (Fig. 5*A*). To establish whether the dispersion of the Hfq foci was a direct response to the presence of N or because of resumption of growth, we repeated the experiment and inoculated bacteria from N-24 either into fresh growth media that was devoid of either N or C, which, could not support the resumption of growth. In media devoid of N (but one that contained C), we failed to detect the dissipation of the Hfq foci even after ~3 h after inoculation (Fig. 5*B*). However, strikingly, the Hfq foci begun to dissipate upon inoculation into media that only contained N but not C (Fig. 5*C*). Thus, we conclude that the Hfq foci are reversible and that their formation and dissipation are a direct response to the availability of N. Further, the reversibility of the Hfq foci also serves to alleviate concerns that the formation of the Hfq foci are due to the inherent tendency of some mCherry tagged proteins to aberrantly aggregate (35).

**Fig. 5.**
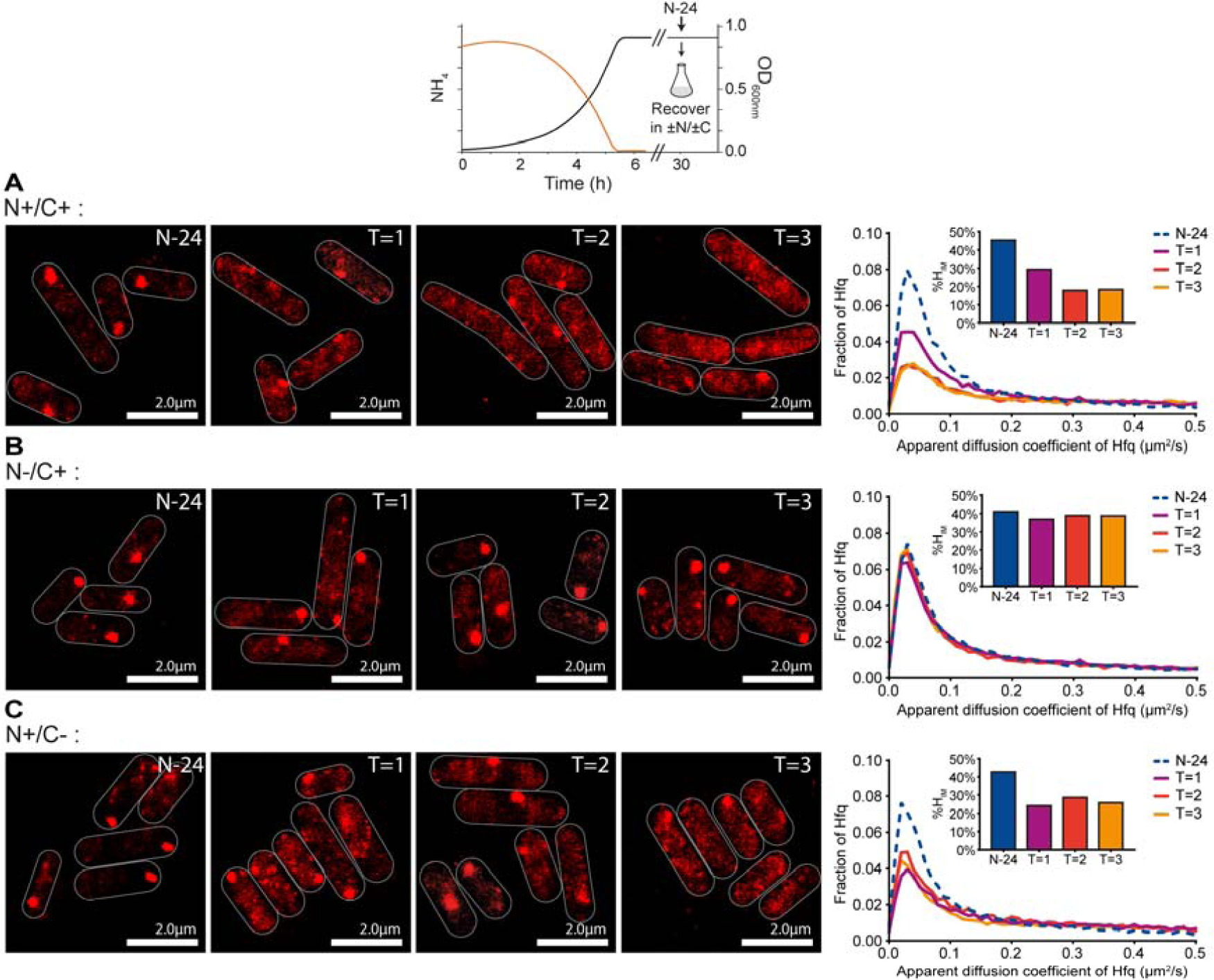
Hfq foci are reversible. Representative PALM images of Hfq from N-24 *E. coli* cells resuspended in fresh media with different combinations of N and C ((*A*) N+/C+, (*B*) N-/C+, (*C*) N+/C-). Images taken hourly post resuspension. The graphs show the distribution of apparent diffusion coefficient of Hfq molecules at indicated time points with corresponding %H_IM_ values shown in inset graphs.

### Hfq foci formation occurs independently of the Ntr response in long-term N starved *E. coli*

The results thus far suggest that foci formation by Hfq is involved in adapting to long-time N starvation. We next wanted to find out whether Hfq foci formation was part of the NtrC activated Ntr response, which is the initial major response to N deficiency. As the Ntr response primarily manifests in global changes in gene expression in response to N starvation, we investigated whether perturbing the transcriptional and translational response with the RNA polymerase inhibitor rifampicin and 50S ribosomal protein inhibitor chloramphenicol, respectively, affected Hfq foci formation. As shown in Fig. 6*A* and 6*B*, the addition of 100 μg/ml rifampicin or 150 μg/ml chloramphenicol to bacteria prior to N- (ND; Fig. 1*A*) or 1 h after N- (N-1) did not markedly affect Hfq foci formation following long-term N starvation (i.e. at N-24) compared to untreated cells. The observation that Hfq foci still form 24 h following N starvation when transcription or translation is perturbed at ND suggests that Hfq foci formation occurs independently of (and subsequently to) the Ntr response. Consistent with this view, the observation that Hfq foci still form 24 h following N starvation when transcription or translation is perturbed at N-1, i.e. once N has run out and bacterial growth has attenuated as a consequence, suggests that Hfq foci formation does not depend on *de novo* transcription or translation once N starvation has set in. Based on these two observations, we expected that Hfq foci formation would be unaffected when the Ntr response is specifically perturbed. Hence, we measured the dynamics of Hfq foci formation in mutant bacteria devoid of NtrC (Δ*glnG*) – the master transcriptional regulator of the Ntr response. As previously reported, the absence of NtrC does prevent *E. coli* to grow under our experimental conditions, although mutant bacteria grew with a moderately slower doubling time than WT bacteria (1). As shown in Fig. 6*D*, Hfq foci were detected in Δ*glnG* bacteria in response to N starvation at N-24, although, the Hfq foci appeared to form moderately faster in the mutant bacteria compared to WT bacteria (Fig. 6*C*; also see Fig. S11). The absence of NtrC also did not affect the ability of the Hfq foci to dissipate when the N starvation stress was alleviated (Fig. 6*D*). Further, as shown in Fig. 6*E* and Fig. 6*F*, respectively, the absence of downstream effectors of the Ntr response, RelA or RpoS, also did not markedly affect Hfq foci formation at N-24, (though foci formation occurred moderately slower in mutant bacterial compared to WT bacteria (Fig. S11) or dissipation when the N starvation stress was alleviated. We also note that the Hfq foci in some Δ*relA* and Δ*rpoS* bacterial cells at N-24 were either absent or irregularly shaped compared to WT bacteria (compare Fig. 6*C* with Fig. 6*E* and 6*F*; also see Fig. S11). Collectively, the results demonstrate that Hfq foci formation is a response to long-term N starvation that occurs independently of the Ntr response and *de novo* gene expression once N starvation has set in.

**Fig. 6.**
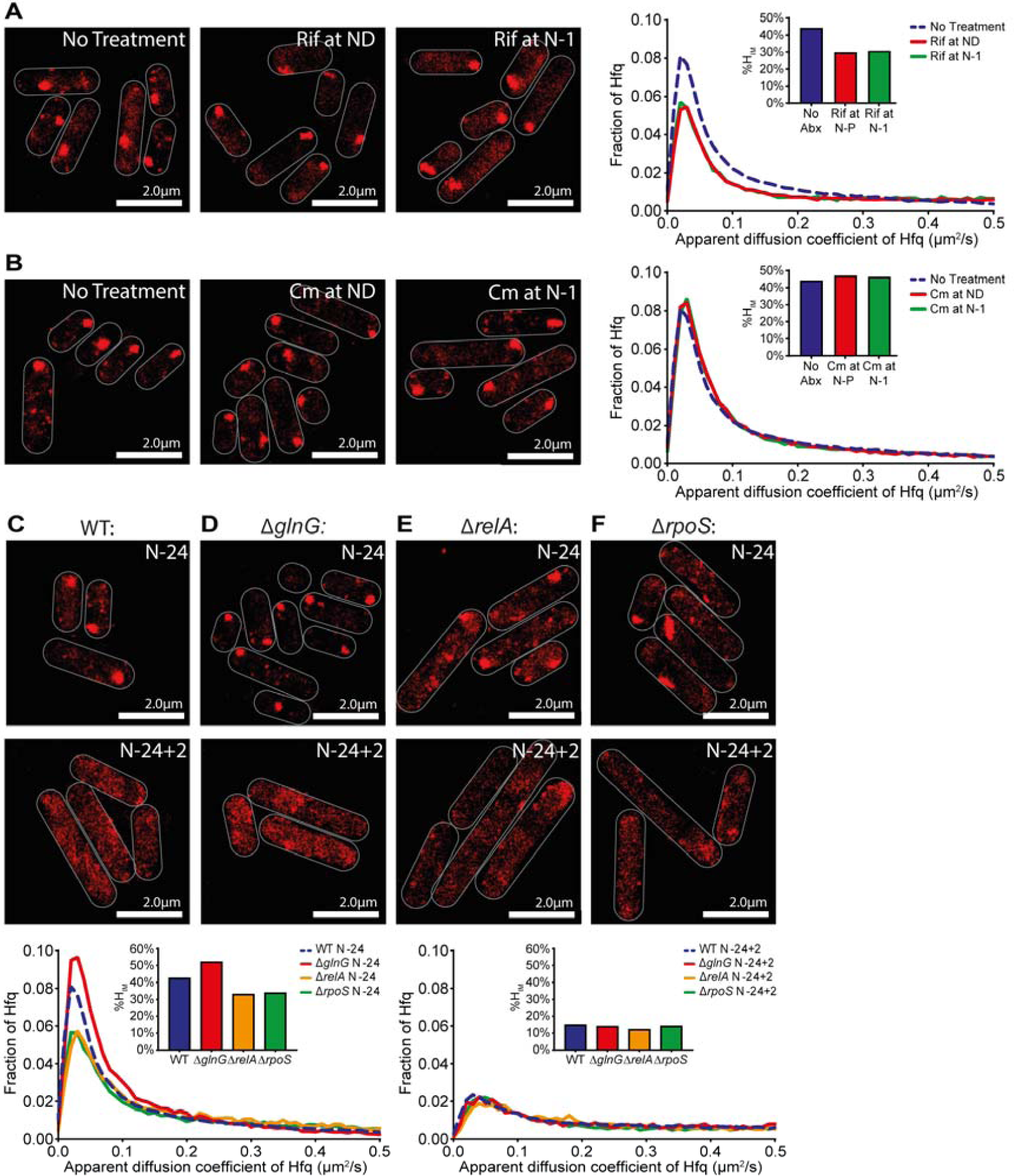
Hfq foci formation occurs independently of the Ntr response in long-term N starved *E. coli*. (*A*) Representative PALM images of Hfq in *E. coli* cells treated with rifampicin (100 μg/ml) at ND (*middle panel*) and 1 h following onset of N starvation (N-1) (*right panel*) and imaged at N-24. The untreated bacteria from N-24 are shown for comparison (*left panel*). Inset graph shows the distribution of apparent diffusion coefficient of Hfq molecules and corresponding %H_IM_ values. (*B*) As in (A) but 150 μg/ml of chloramphenicol was used. (*C*)-(*F*) Representative PALM images of Hfq in *E. coli* cells in WT (*C*), Δ*glnG* (*D*), Δ*relA* (*E*) and Δ*rpoS* (*F*) strains at N-24 and 2h (N-24+2) following alleviation of N starvation stress (as in Fig. 5*A*). Graphs showing the distribution of apparent diffusion coefficient of Hfq molecules and the %H_IM_ values at the sampled time points.

## Conclusion

Although the adaptive response to N starvation in *E. coli* has been well investigated, many of these studies have been conducted with bacteria subjected to short-term N starvation (e.g. (4, 19, 36–38)). This study has provided insights into how the adaptive response to N starvation can be subjected to temporal coordination as N starvation ensues for a long time and has assigned a critical role for the RNA binding protein Hfq in this process. This study has uncovered that, as N starvation ensues, Hfq forms a single and reversible focus-like structure in long-term N starved *E. coli* cells, which is markedly distinct from clustering of Hfq molecules seen previously in *E. coli* under different growth or stress conditions (18, 31, 32). Significantly, the formation of the Hfq foci occurs independently of (i) the NtrC activated Ntr response, which occurs at the onset of N starvation and (ii) *de novo* gene expression in bacteria experiencing N starvation. As Hfq is a global regulator of bacterial cell function, it is difficult to unambiguously separate, at this stage in our analyses, the other functions of Hfq from its specific role in the adaptive response to long-term N starvation in mechanistic detail. Although future work will now focus on defining the composition and organisation of the Hfq foci, the experiments with T7 unambiguously indicate that the Hfq foci contribute to adapting *E. coli* cell function as N starvation ensues. We speculate that the Hfq foci could be a ribonucleoprotein complex which degrade unwanted RNA molecules to release N from the nitrogenous RNA molecules or sequester cellular resources to maximise the chances of survival as N starvation conditions ensue. Thus, the absence of Hfq foci could result in the ‘mismanagement’ of cellular processes as N starvation ensues. Such a scenario would explain the inability of T7 phage, which heavily relies on host bacterial resources, to replicate in N-24 Δ*hfq* bacteria where the Hfq foci cannot form. It is thus tempting to speculate that the Hfq foci resemble ribonucleoprotein complexes observed in stressed eukaryotic cells, which are involved in broad aspects of managing RNA metabolism as direct response to stress (39–41). Indeed, the gradual accumulation of the Hfq foci as a function of time under N stress, their size and their dissipation upon alleviation of the N stress are some of the properties Hfq foci share with ribonucleoprotein complexes observed in stressed eukaryotic cells.

In summary, a recent study by Kannaiah et al (18) and this study have now independently shown that Hfq has the propensity to assemble into foci like structures in *E. coli* experiencing stress. In case of the former study, the Hfq foci formed to locate small RNAs at the poles of the cells as a mechanism of spatiotemporal regulation of gene expression in response to envelope stress. The spatiotemporal regulation of bacterial processes is an emerging area of research in bacteriology and the Hfq foci described in this study represents a mechanism to spatiotemporally manage cell function during long-term N starvation. Importantly, this study has also indicated that adaptive processes in nutrient starved and growth attenuated bacteria can occur independently of *de novo* gene expression and major regulatory pathways that have evolved to allow bacteria adapt to nutrient stresses.

## Material and Methods

### Bacterial strains and plasmids

All strains used in this study were derived from *Escherichia coli* K-12 and are listed in Supplementary Table 1. The Hfq-PAmCherry and MetJ-PAmCherry strains were constructed using the λ Red recombination method (42) to create an in-frame fusion encoding a linker sequence and PAmCherry, followed by a kanamycin resistance cassette (amplified from the KF26 strain (43)) to the 3′ end of *hfq* and *metJ*. The ProQ-PAmCherry reporter strain (Δ*proQ*+pACYC*-proQ-PAmCherry*) was constructed using a Δ*proQ* strain (Provided by Prof. Jörg Vogel, University of Würzburg). The pACYC-ProQ-PAmCherry plasmid was made by Gibson assembly and used to express ProQ-PAmCherry under the native promoter of *proQ* (44). The Hfq_Δ73-102_–PAmCherry strain was made using the λ Red recombination method (42), similar to construction of the Hfq-PAmCherry strain, but with an in-frame fusion of the linker-PAmCherry sequence to replace amino acids 73-102. Gene deletions (Δ*glnG*, Δ*relA* or Δ*rpoS*) were introduced into the Hfq-PAmCherry strain as described previously (19): Briefly, the knockout alleles were transduced using the P1*vir* bacteriophage with strains from the Keio collection (45) serving as donors.

### Bacterial growth conditions

Bacteria were grown in Gutnick minimal medium (33.8□mM KH_2_PO_4_, 77.5□mM K_2_HPO_4_, 5.74□mM K_2_SO_4_, 0.41□mM MgSO_4_) supplemented with Ho-LE trace elements (46), 0.4□% (w/v) glucose as the sole C source and NH_4_Cl as the sole N source. Overnight cultures were grown at 37□°C, 180□r.p.m. in Gutnick minimal medium containing 10□mM NH_4_Cl. For the N starvation experiments, 3□mM NH_4_Cl was used (see text for details). For C starvation experiments, bacteria were grown in Gutnick minimal medium containing 0.06% (w/v) glucose and 10 mM NH_4_Cl. The NH_4_Cl concentrations in the growth media were determined using the Aquaquant ammonium quantification kit (Merck Millipore, UK) as per the manufacturer’s instructions. The proportion of viable cells in the bacterial population was determined by measuring CFU/ml from serial dilutions on lysogeny broth agar plates. Complementation experiments used pBAD24-*hfq*-3xFLAG, or pBAD18 as the empty vector control, in Gutnick minimal medium supplemented with 0.2% (w/v) L-arabinose at t=0, for induction of gene expression. To observe Hfq foci dissipation, 25 ml of N-24 culture was centrifuged at 3,200 *g* and resuspend in fresh Gutnick minimal medium containing different combinations of 0.4% (w/v) glucose and 3 mM NH_4_Cl (see text for details).

### Photoactivated localization microscopy (PALM) and single molecule tracking (SMT)

For the PALM and SMT experiments, the Hfq-PAmCherry (and derivatives), KF26, MetJ-PAmCherry and ProQ-PAmCherry reporter strains were used. The bacterial cultures were grown as described above and samples were taken at the indicated time points, then imaged and analysed as previously described (43, 47). Briefly, 1 ml of culture was centrifuged, washed and resuspended in a small amount of Gutnick minimal medium without any NH_4_Cl + 0.4% glucose; samples taken at N+ were resuspended in Gutnick minimal medium with 3 mM NH_4_Cl + 0.4% glucose. For the C starvation experiments, C- and C-24 samples were resuspended in Gutnick minimal medium with 10 mM NH_4_Cl but no glucose; C+ samples were resuspended in Gutnick minimal medium with 10 mM NH_4_Cl and 0.06% glucose. One μl of the resuspended culture was then placed on a Gutnick minimal medium agarose pad (1x Gutnick minimal medium with no NH_4_Cl + 0.4% glucose with 1% (w/v) agarose); samples taken at N+ were placed on a pad made with Gutnick minimal medium with 3 mM NH_4_Cl. For the C starvation experiments, C- and C-24 samples were placed on a pad containing 10 mM NH_4_Cl but no glucose; C+ samples were placed on a pad containing 10 mM NH_4_Cl and 0.06% glucose. Cells were imaged on a PALM-optimized Nanoimager (Oxford Nanoimaging, www.oxfordni.com) with 15 millisecond exposures, at 66 frames per second over 10,000 frames. Photoactivatable molecules were activated using 405 nm and 561 nm lasers. For SMT, the Nanoimager software was used to localize the molecules by fitting detectable spots of high photon intensity to a Gaussian function. The Nanoimager software SMT function was then used to track individual molecules and draw trajectories of individual molecules over multiple frames, using a maximum step distance between frames of 0.6 μm and a nearest-neighbour exclusion radius of 0.9 μm. The software then calculated the apparent diffusion coefficients (*D\**) for each trajectory over at least four steps, based on the mean squared displacement of the molecule.

### Immunoblotting

Immunoblotting was conducted in accordance to standard laboratory protocols (48). The following antibodies were used: mouse monoclonal anti-DnaK 4RA2 at 1□:□10,□000 dilution (Enzo, ADI-SPA-880) and HRP Goat anti-mouse IgG (BioLegend, 405306) at 1□:□10,□000 dilution. ECL Prime Western blotting detection reagent (GE Healthcare, RPN2232) was used to develop the blots, which were analysed on the ChemiDoc MP imaging system and bands quantified using Image Lab software.

### Purification of insoluble protein fractions

Insoluble protein fractions were purified exactly as previously described (30). Briefly, aliquots of bacterial cultures (5-20 ml) were cooled on ice and cell pellets harvested by centrifugation at 3,200 *g* for 10 min at 4 °C. To extract the insoluble protein fraction, pellets were resuspended in 40 μl Buffer A (10 mM Potassium phosphate buffer, pH 6.5, 1 mM EDTA, 20% (w/v) sucrose, 1 mg/ml lysozyme) and incubated on ice for 30 min. Cells were lysed by adding 360 μl of Buffer B (10 mM Potassium phosphate buffer, pH 6.5, 1 mM EDTA) followed by sonication (10 s pulse on, 10 s off, 40% amplitude, repeated x 6 (2 min total time)) on ice. Intact cells were removed by centrifugation at 2,000 *g* for 15 min at 4 °C.

The insoluble fraction was collected by centrifugation at 15,000 *g* for 20 min at 4 °C and kept at −80 °C. Pellets were resuspended in 400 μl of Buffer B and centrifuged at 15,000 *g* for 20 min at 4 °C. The pellet was then resuspended in 320 μl Buffer B plus 80 μl of 10% (v/v) NP40. Aggregated proteins were then collected by centrifugation at 15,000 *g* for 30 min at 4 °C. This wash step was repeated. The NP40 insoluble pellets were washed with 400 μl Buffer B. The final insoluble protein pellet was resuspended in 200 μl of Buffer B and 200 μl LDS loading dye and samples were run on a 12 % (w/v) sodium dodecyl sulphate polyacrylamide gel and analysed by Coomassie stain and immunoblotting, in accordance to standard laboratory protocols (48). To obtain whole cell extracts, bacterial pellets were resuspended in 40 µl of Buffer A (with no lysozyme) and 50 µl of 2x lithium dodecyl sulphate loading dye was added before boiling the sample for 5 min and analysis using sodium dodecyl sulphate polyacrylamide gel separation as above.

### T7 phage infection assay

Bacterial cultures were grown in Gutnick minimal media as described above to the indicated time points. Samples were taken and diluted to OD_600nm_ of 0.3 in Gutnick minimal media containing ~3 mM NH_4_Cl to a final volume of 500 μl and transferred to a flat-bottom 48-well plate together with T7 phage at a final concentration of 4.2×10^9^ phage/ml. Cultures were then grown at 37 °C, shaking at 700 r.p.m. in a SPECTROstar Nano Microplate Reader (BMG LABTECH) and OD_600nm_ readings were taken every 10 mins.

## Acknowledgments

This work was supported by Wellcome Trust Investigator Award 100958 to S.W. and a Medical Research Council Ph.D. studentship to J.M.

## Figures and Tables

**Fig. S1.**
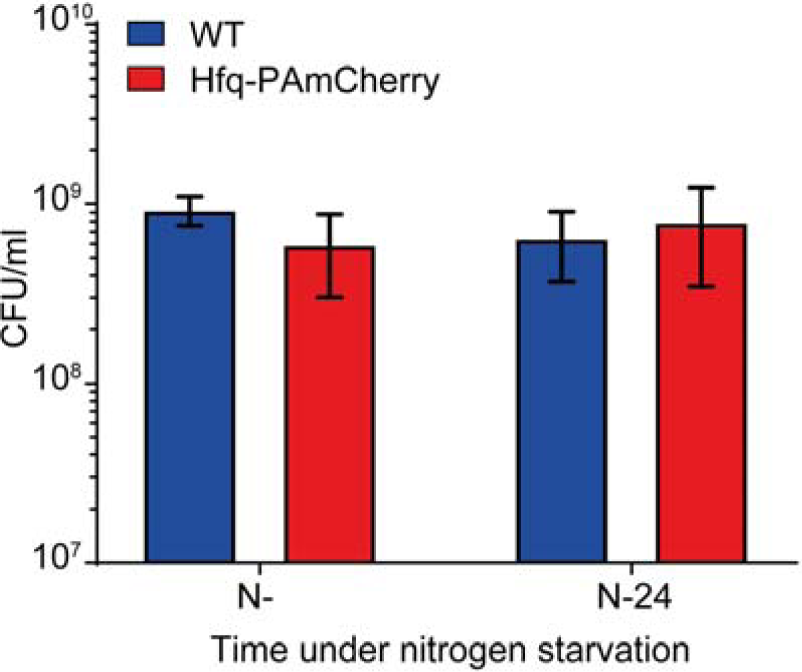
The PAmCherry tag on Hfq does not affect cell viability. Viability of WT and Hfq-PAmCherry *E. coli* at N- and N-24 measured by CFU counts. Error bars represent standard deviation (n□ =□3).

**Fig. S2.**
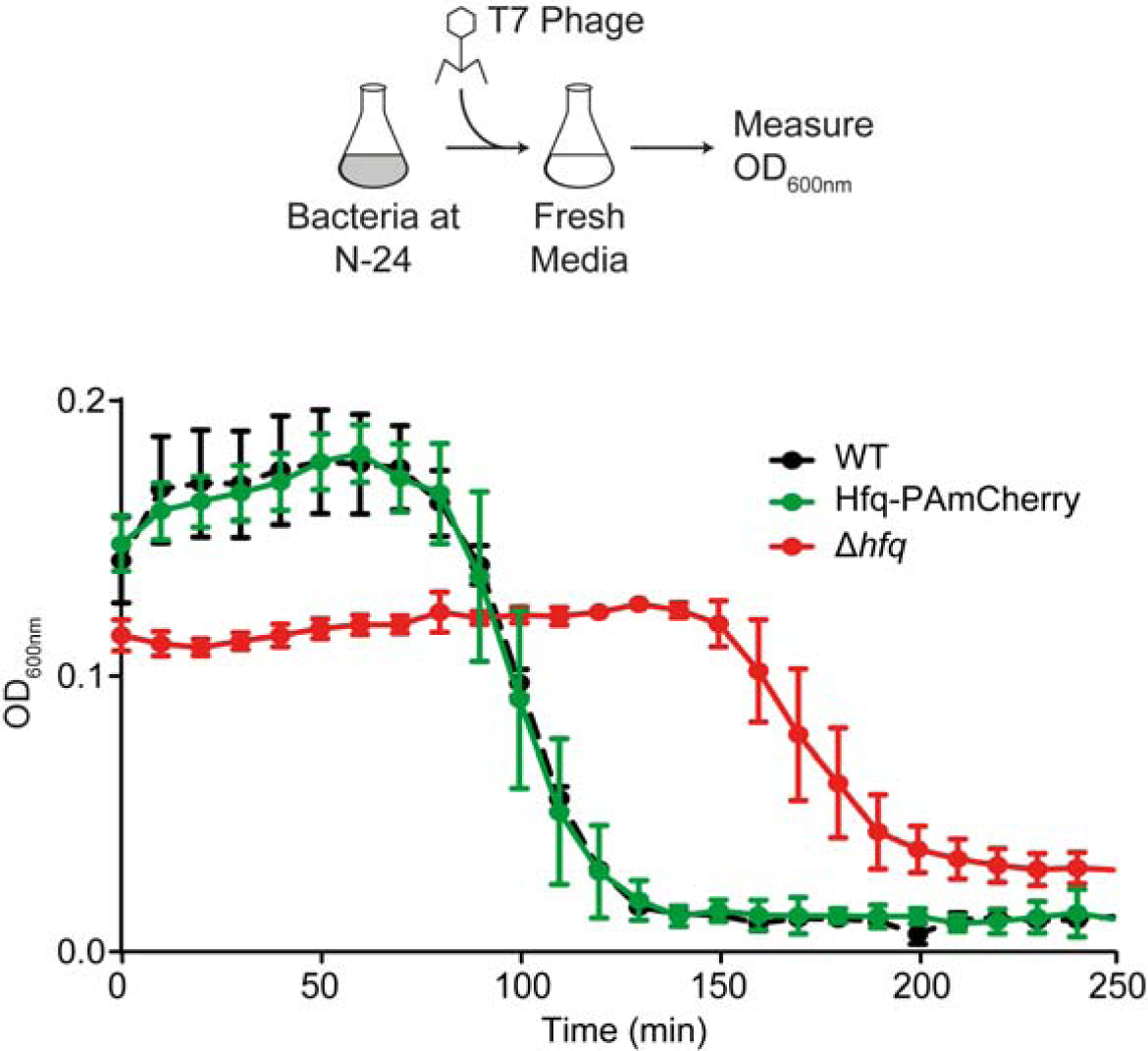
The PAmCherry tag on Hfq does not affect the ability for T7 phage to replicate *E. coli* cells. Graph showing the optical density as a function of time of WT, Hfq-PAmCherry and Δ*hfq E. coli* cells from N-24 following infection with T7 phage.

**Fig. S3.**
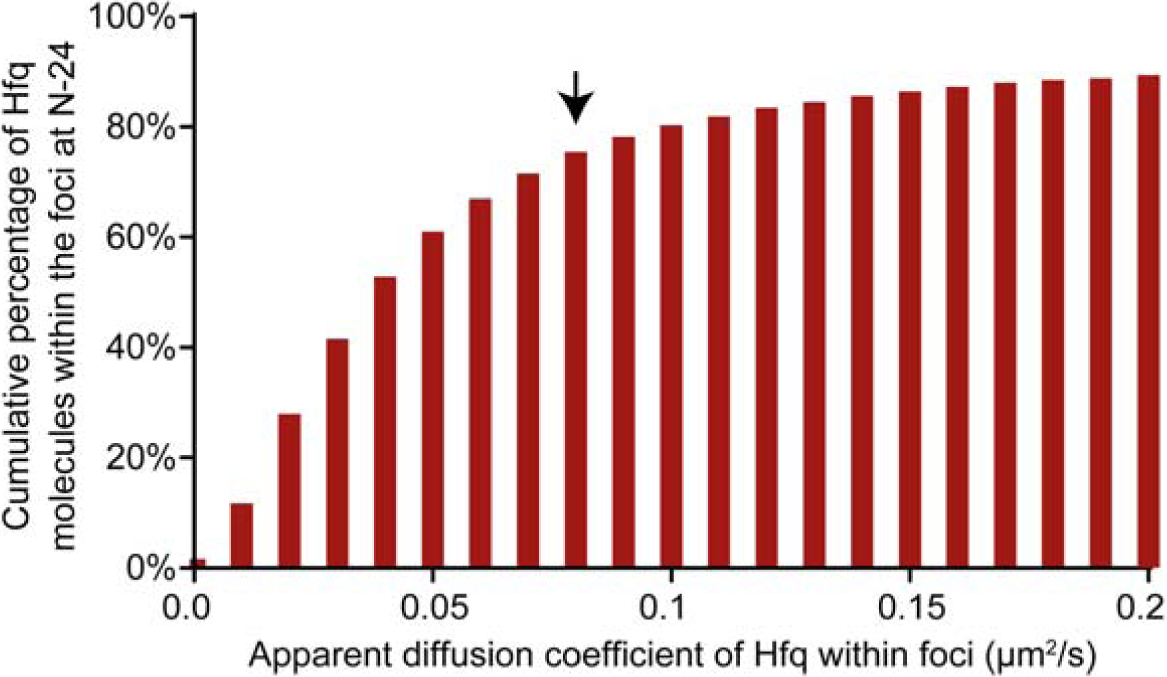
The majority of Hfq molecules in the foci have an apparent diffusion coefficient of <0.08. Graph showing the cumulative proportion of Hfq molecules within the foci with increasing *D\** values. Arrow indicates cut-off value used for defining %H_IM_. See text for details.

**Fig. S4.**
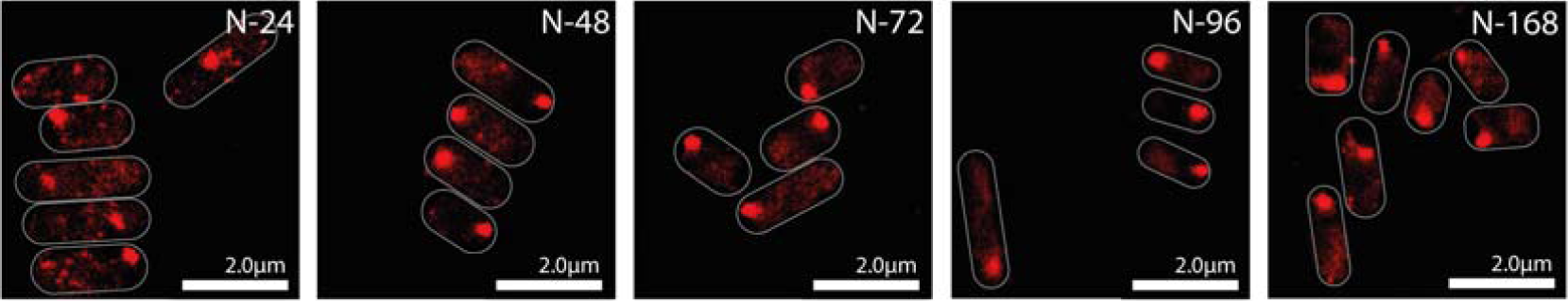
Hfq foci persist for at least a 168 h under N starvation. Representative PALM images of Hfq in *E. coli* cells under long-term N starvation. Images taken at indicated time points.

**Fig. S5.**
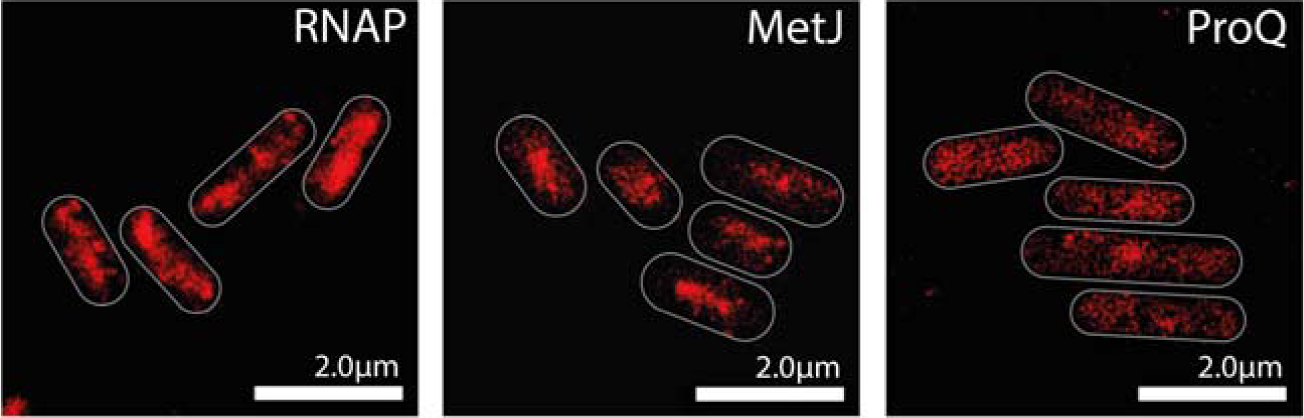
Hfq foci formation is not a generic feature of proteins during long-term N starvation. Representative PALM images of PAmCherry tagged: RNA Polymerase (RNAP) (*left panel*); MetJ (*middle panel*); and ProQ (*right panel*) in *E. coli* cells at N-24.

**Fig. S6.**
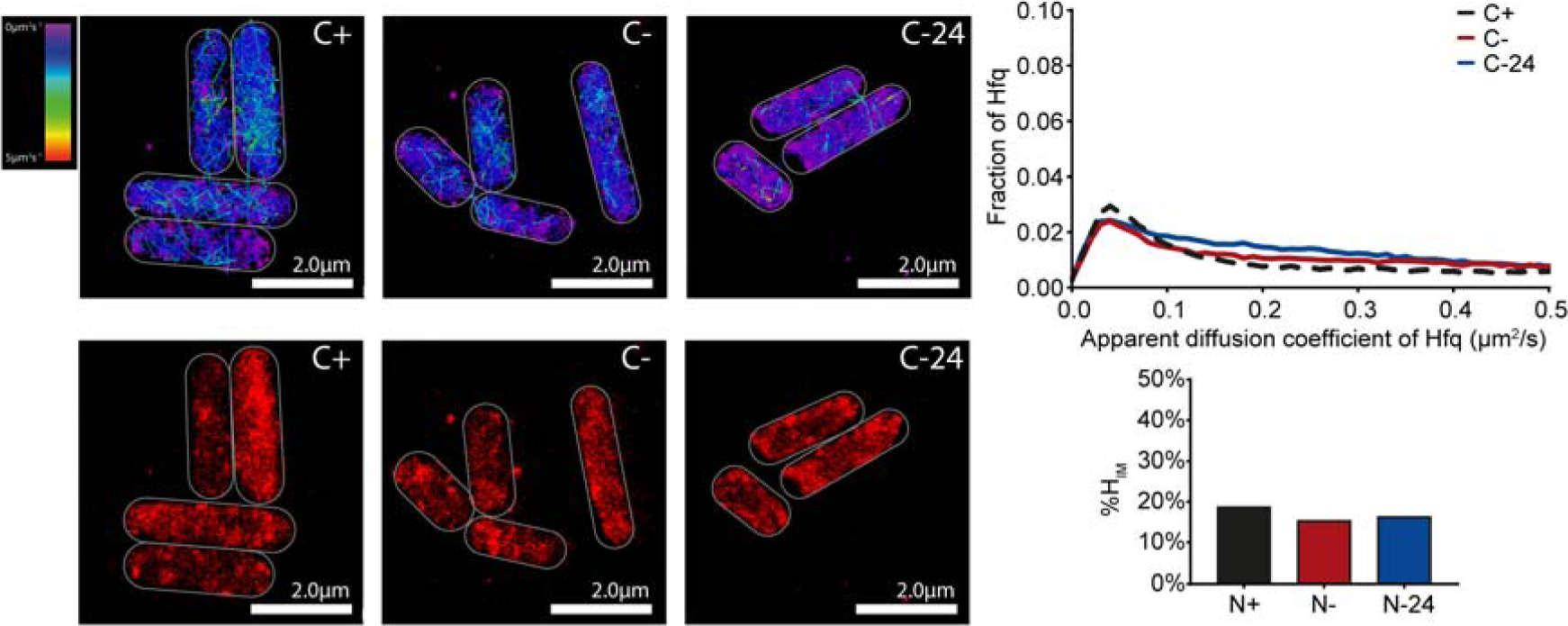
Hfq does not form foci in long-term C starved *E. coli* cells. Representative single molecule tracks (*top panel*) and PALM images (*bottom panel*) of Hfq in *E. coli* cells from C+, C- and C-24. The graph shows the distribution of apparent diffusion coefficient of Hfq molecules at indicated time points and corresponding %H_IM_ values shown in the inset graph.

**Fig. S7.**
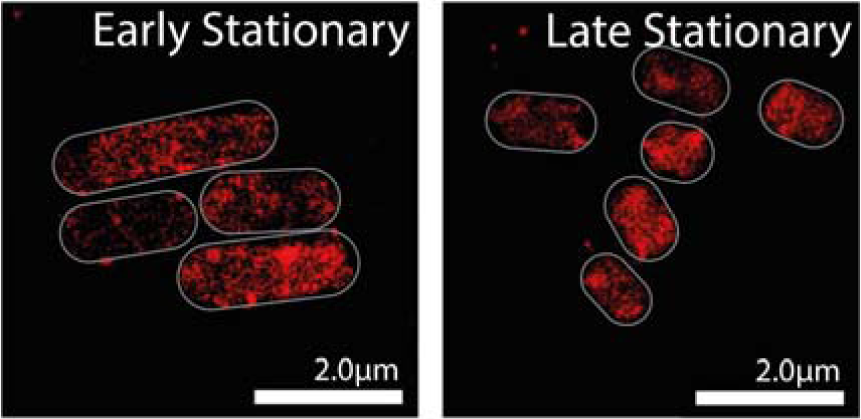
Hfq does not forms foci during stationary phase in lysogeny broth. Representative PALM images of Hfq in *E. coli* cells grown to early (1 h) and late (24 h) stationary phase in lysogeny broth.

**Fig. S8.**
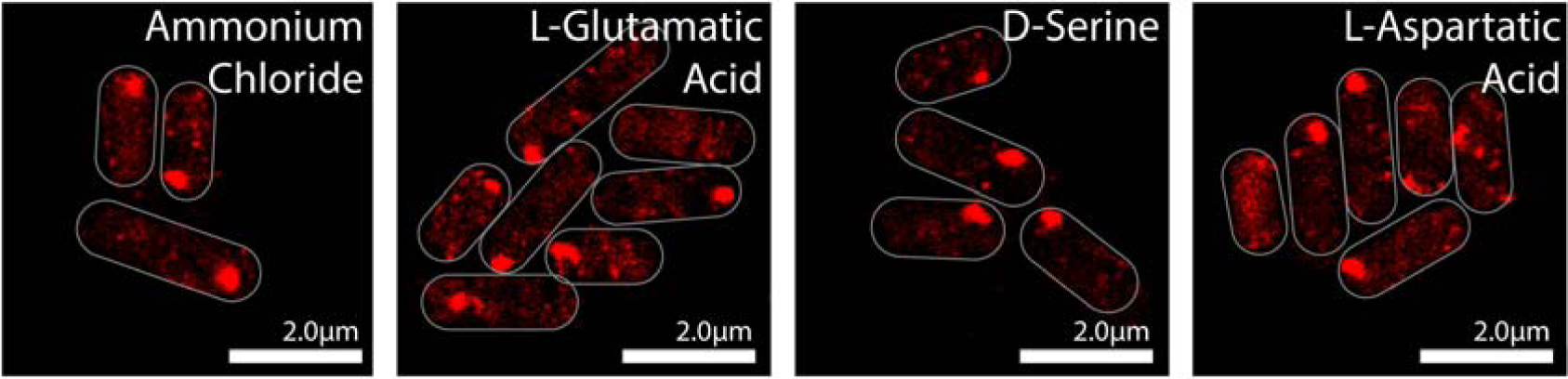
Hfq foci form in response to starvation of diverse N sources. Representative PALM images of Hfq in *E. coli* cells when 3mM of NH_4_Cl (*first panel*), L-Glutamatic acid (*second panel*), D-Serine (*third panel*) or L-Aspartatic acid (*fourth panel*) was used as the sole N source. Cells for imaging were sampled following 24 h of growth, using growth in NH_4_Cl as the reference time point.

**Fig. S9.**
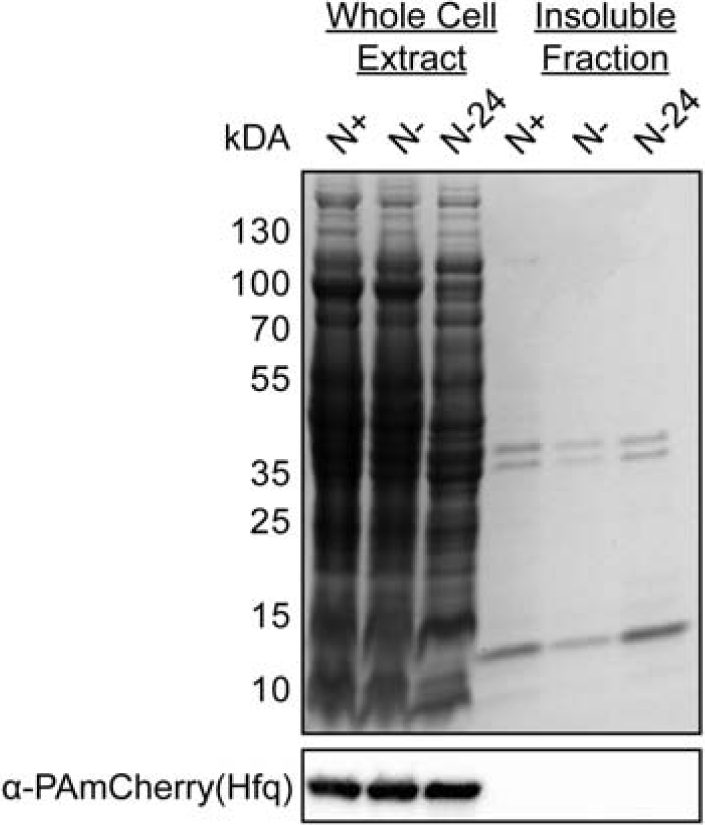
Hfq foci are not aberrant aggregates of Hfq molecules in long-term N starved *E. coli.* Representative Coomassie stained denaturing gel of whole cell extracts and insoluble protein fraction of *E. coli* cells containing Hfq-PAmCherry from N+, N- and N-24 (*top panel*) and immunoblot of the section containing Hfq (*bottom panel*) using antibodies against the PAmCherry tag.

**Fig. S10.**
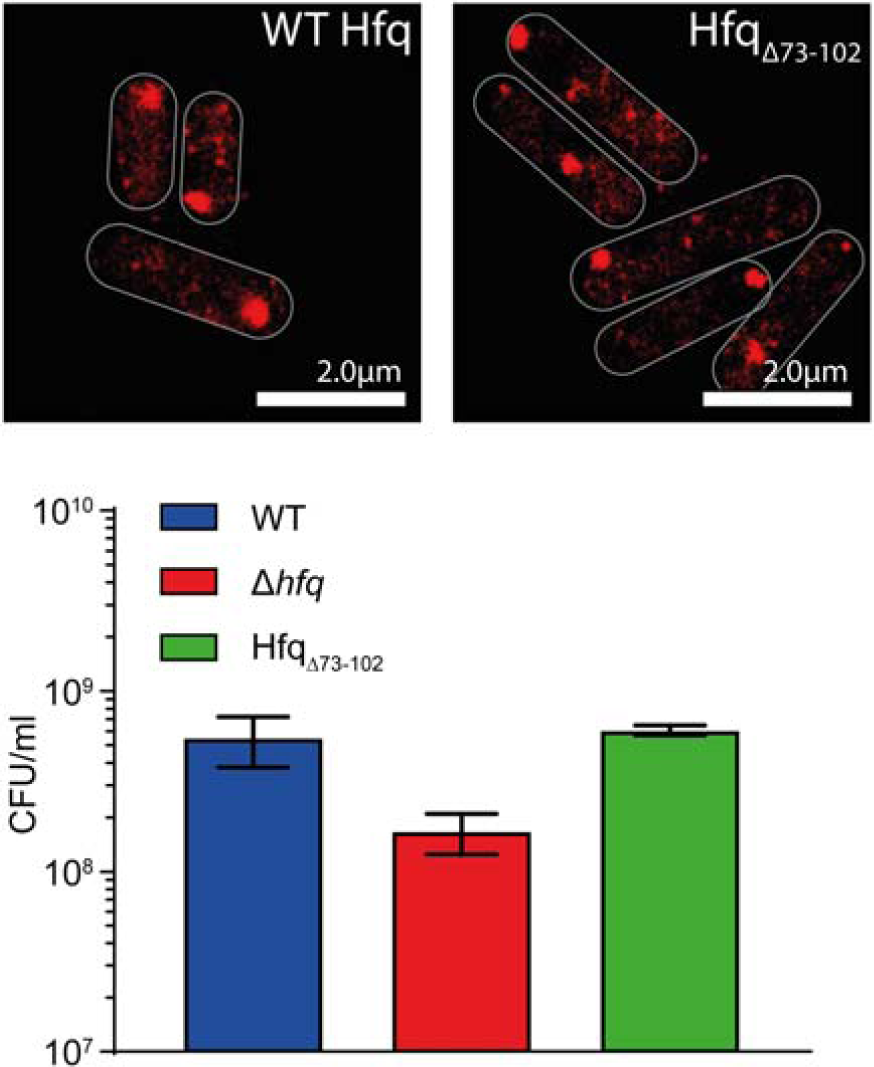
Hfq foci formation is not dependent on the C-terminal amino acid residues of Hfq. Representative PALM images of full length Hfq (*left panel*) and C-terminally truncated Hfq (Hfq_Δ73-102_) (*right panel*) in *E. coli* cells at N-24. The graph shows the viability of WT Δ*hfq* and Hfq_Δ73-102_ *E. coli* at N-24 measured by CFU counts. Error bars represent standard deviation (n□= □3).

**Fig. S11.**
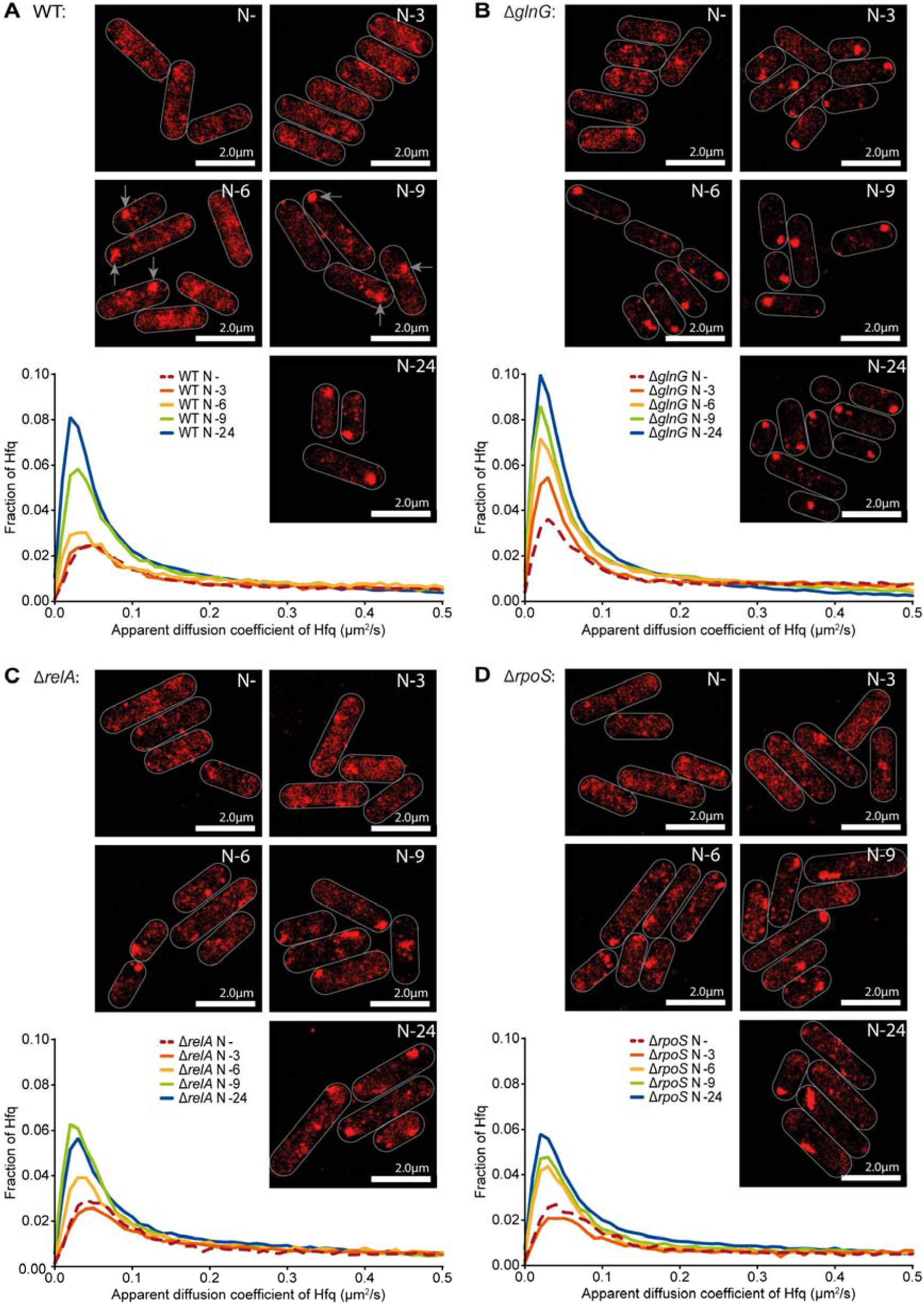
Hfq foci formation occurs independently of the Ntr response in long-term N starved *E. coli*. Representative PALM images of Hfq in *E. coli* cells in (*A*) WT, (*B*) Δ*glnG*, (*C*) Δ*relA* and (*D*) Δ*rpoS* strains, as a function of time under N starvation. The graph shows the distribution of apparent diffusion coefficient of Hfq molecules at the indicated sampling time points.

**Table S1.** *E. coli* strains and plasmids used in this study.

| Strains |  |  |
| --- | --- | --- |
| Name | Description | Source or Reference |
| Wild-type (BW25113) | *E. coli* K-12 (*araD-araB*)567 $\Delta$(*rhaD-rhaB*)<br>568 Δ*lacZ*4787 (*::rrnB-3*) *hsdR*514 *rph*-1 | *E. coli* Genetic Stock Center |
| Δ*hfq* (JW4130) | BW25113 Δ*hfq::kan* | *E. coli* Genetic Stock Center |
| Δ*rpoS* (JW5437) | BW25113 Δ*rpoS::kan* | *E. coli* Genetic Stock Center |
| Wild-type (MG1655) | *E. coli* K-12 *rph*-1 | *E. coli* Genetic Stock Center |
| Δ*hfq* (MG1655) | MG1655 Δ*hfq::kan* | |
| Hfq-PAmCherry | MG1655 *hfq*-PAmCherry-*kan* | This Study |
| MetJ-PAmCherry | MG1655 *metJ*-PAmCherry-*kan* | This Study |
| RpoC-PAmCherry (KF26) | MG1655 *rpoC*-PAmCherry-*kan* | (43) |
| Δ*ProQ* | MG1655 Δ*proQ::kan* | Provided by Prof. Jörg Vogel, University of Würzburg |
| HfqΔ<sub>73-102</sub>-PAmCherry | MG1655 *hfq*Δ<sub>73-102</sub>-PAmCherry-*kan* | This Study |
| Δ*glnG* Hfq-PAmCherry | Hfq-PAmCherry Δ*glnG::kan* | This Study |
| Δ*relA* Hfq-PAmCherry | Hfq-PAmCherry Δ*relA::kan* | This Study |
| Δ*rpoS* Hfq-PAmCherry | Hfq-PAmCherry Δ*rpoS::kan* | This Study |
| Plasmids |  |  |
| Name | Description | Source/Reference |
| pBAD24-*hfq*-FLAG | pBAD24 expressing *hfq*-3xFLAG under an arabinose inducible promoter | Provided by Prof. Jörg Vogel, University of Würzburg |
| pBAD18 | Empty pBAD18 | (49) |
| pACYC-*proQ*-PAmCherry | Modified pACYC backbone (-TcR, +MCS) expressing *proQ*-PAmCherry under the native *proQ* promoter | This Study |

**Table S2.**
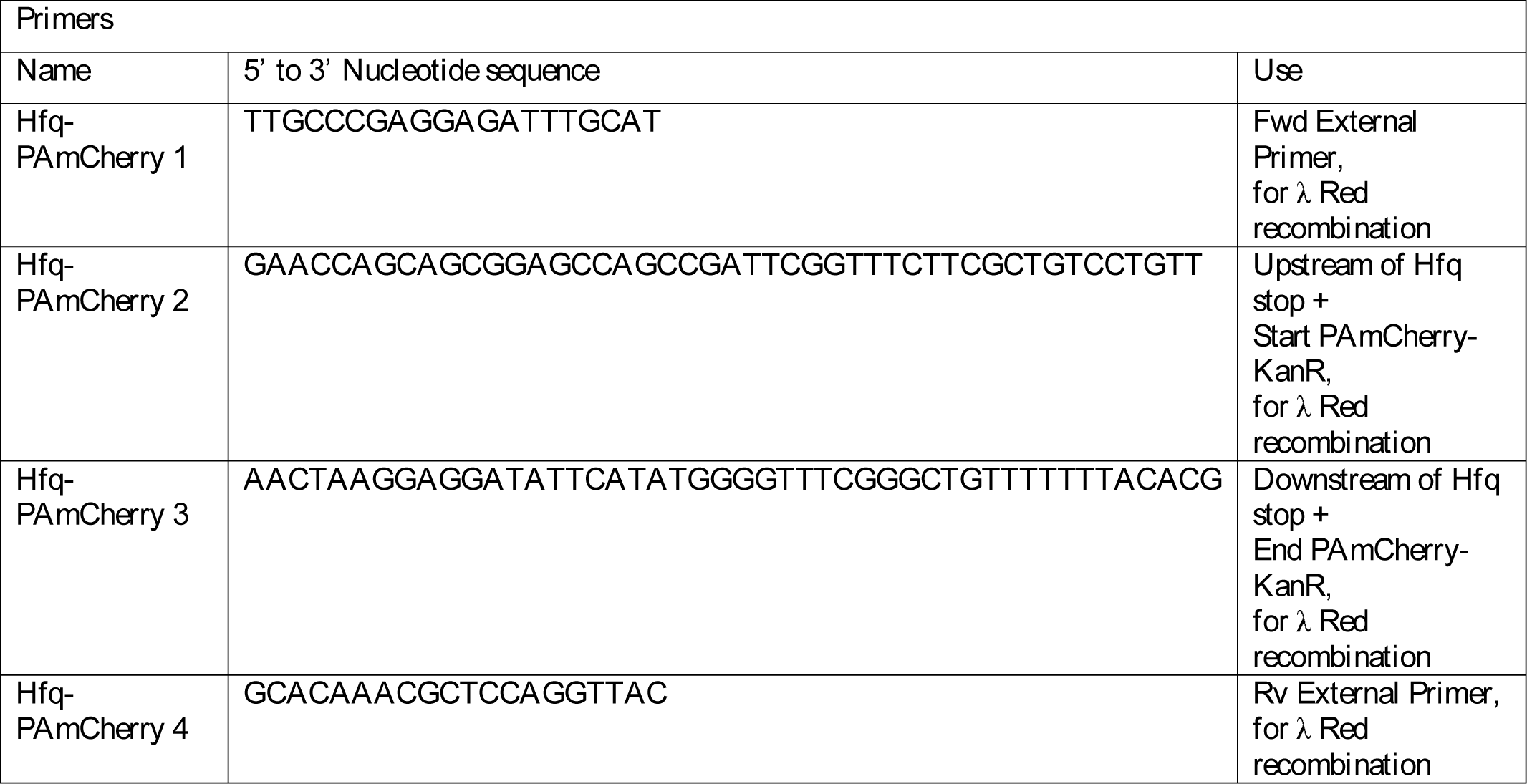
Primers used in this study.

## References

1. Brown DR, Barton G, Pan Z, Buck M, & Wigneshweraraj S (2014) Combinatorial stress responses: direct coupling of two major stress responses in Escherichia coli. Microb Cell 1(9):315–317.

2. Switzer A, Brown DR, & Wigneshweraraj S (2018) New insights into the adaptive transcriptional response to nitrogen starvation in Escherichia coli. Biochem Soc Trans 46(6):1721–1728.

3. Gyaneshwar P, et al. (2005) Sulfur and nitrogen limitation in Escherichia coli K-12: specific homeostatic responses. J Bacteriol 187(3):1074–1090.

4. Zimmer DP, et al. (2000) Nitrogen regulatory protein C-controlled genes of Escherichia coli: scavenging as a defense against nitrogen limitation. Proc Natl Acad Sci U S A 97(26):14674–14679.

5. Amin SV, et al. (2016) Novel small RNA (sRNA) landscape of the starvation-stress response transcriptome of Salmonella enterica serovar typhimurium. RNA Biol 13(3):331–342.

6. Holmqvist E & Wagner EGH (2017) Impact of bacterial sRNAs in stress responses. Biochem Soc Trans 45(6):1203–1212.

7. Kwenda S, et al. (2016) Discovery and profiling of small RNAs responsive to stress conditions in the plant pathogen Pectobacterium atrosepticum. BMC Genomics 17:47.

8. Gerrick ER, et al. (2018) Small RNA profiling in Mycobacterium tuberculosis identifies MrsI as necessary for an anticipatory iron sparing response. Proc Natl Acad Sci U S A 115(25):6464–6469.

9. Mohd-Padil H, et al. (2017) Identification of sRNA mediated responses to nutrient depletion in Burkholderia pseudomallei. Sci Rep 7(1):17173.

10. Balasubramanian D & Vanderpool CK (2013) New developments in post-transcriptional regulation of operons by small RNAs. RNA Biol 10(3):337–341.

11. Desnoyers G, Bouchard MP, & Masse E (2013) New insights into small RNA-dependent translational regulation in prokaryotes. Trends Genet 29(2):92–98.

12. Andrade JM, Dos Santos RF, Chelysheva I, Ignatova Z, & Arraiano CM (2018) The RNA-binding protein Hfq is important for ribosome biogenesis and affects translation fidelity. EMBO J 37(11).

13. Lee T & Feig AL (2008) The RNA binding protein Hfq interacts specifically with tRNAs. RNA 14(3):514–523.

14. Andrade JM, Pobre V, Matos AM, & Arraiano CM (2012) The crucial role of PNPase in the degradation of small RNAs that are not associated with Hfq. RNA 18(4):844–855.

15. Mohanty BK, Maples VF, & Kushner SR (2004) The Sm-like protein Hfq regulates polyadenylation dependent mRNA decay in Escherichia coli. Mol Microbiol 54(4):905–920.

16. Le Derout J, et al. (2003) Hfq affects the length and the frequency of short oligo(A) tails at the 3’ end of Escherichia coli rpsO mRNAs. Nucleic Acids Res 31(14):4017–4023.

17. Moll I, Leitsch D, Steinhauser T, & Blasi U (2003) RNA chaperone activity of the Sm-like Hfq protein. EMBO Rep 4(3):284–289.

18. Kannaiah S, Livny J, & Amster-Choder O (2019) Spatiotemporal Organization of the E. coli Transcriptome: Translation Independence and Engagement in Regulation. Mol Cell.

19. Brown DR, Barton G, Pan Z, Buck M, & Wigneshweraraj S (2014) Nitrogen stress response and stringent response are coupled in Escherichia coli. Nat Commun 5:4115.

20. Battesti A, Majdalani N, & Gottesman S (2011) The RpoS-mediated general stress response in Escherichia coli. Annu Rev Microbiol 65:189–213.

21. Soper T, Mandin P, Majdalani N, Gottesman S, & Woodson SA (2010) Positive regulation by small RNAs and the role of Hfq. Proc Natl Acad Sci U S A 107(21):9602–9607.

22. Majdalani N, Chen S, Murrow J, St John K, & Gottesman S (2001) Regulation of RpoS by a novel small RNA: the characterization of RprA. Mol Microbiol 39(5):1382–1394.

23. Majdalani N, Cunning C, Sledjeski D, Elliott T, & Gottesman S (1998) DsrA RNA regulates translation of RpoS message by an anti-antisense mechanism, independent of its action as an antisilencer of transcription. Proc Natl Acad Sci U S A 95(21):12462–12467.

24. Tabib-Salazar A, et al. (2018) T7 phage factor required for managing RpoS in Escherichia coli. Proc Natl Acad Sci U S A 115(23):E5353–E5362.

25. Persson F, Linden M, Unoson C, & Elf J (2013) Extracting intracellular diffusive states and transition rates from single-molecule tracking data. Nat Methods 10(3):265–269.

26. Seongjin Park KP, Emily M. Heideman, Marie-Claude Carrier, Matthew A. Reyer, Wei Liu, Eric Massé, View ORCID ProfileJingyi Fei (Dynamic interactions between the RNA chaperone Hfq, small regulatory RNAs and mRNAs in live bacterial cells. https://doi.org/10.1101/2020.01.13.903641.

27. Westermann AJ, et al. (2019) The Major RNA-Binding Protein ProQ Impacts Virulence Gene Expression in Salmonella enterica Serovar Typhimurium. MBio 10(1).

28. Holmqvist E, Li L, Bischler T, Barquist L, & Vogel J (2018) Global Maps of ProQ Binding In Vivo Reveal Target Recognition via RNA Structure and Stability Control at mRNA 3’ Ends. Mol Cell 70(5):971–982 e976.

29. Smirnov A, et al. (2016) Grad-seq guides the discovery of ProQ as a major small RNA-binding protein. Proc Natl Acad Sci U S A 113(41):11591–11596.

30. Tomoyasu T, Mogk A, Langen H, Goloubinoff P, & Bukau B (2001) Genetic dissection of the roles of chaperones and proteases in protein folding and degradation in the Escherichia coli cytosol. Molecular microbiology 40(2):397–413.

31. Fortas E, et al. (2015) New insight into the structure and function of Hfq C-terminus. Biosci Rep 35(2).

32. Taghbalout A, Yang Q, & Arluison V (2014) The Escherichia coli RNA processing and degradation machinery is compartmentalized within an organized cellular network. Biochem J 458(1):11–22.

33. Vecerek B, Rajkowitsch L, Sonnleitner E, Schroeder R, & Blasi U (2008) The C-terminal domain of Escherichia coli Hfq is required for regulation. Nucleic Acids Res 36(1):133–143.

34. Sonnleitner E, et al. (2004) Functional effects of variants of the RNA chaperone Hfq. Biochem Biophys Res Commun 323(3):1017–1023.

35. Landgraf D, Okumus B, Chien P, Baker TA, & Paulsson J (2012) Segregation of molecules at cell division reveals native protein localization. Nat Methods 9(5):480–482.

36. Schumacher J, et al. (2013) Nitrogen and carbon status are integrated at the transcriptional level by the nitrogen regulator NtrC in vivo. MBio 4(6):e00881–00813.

37. Bren A, et al. (2016) Glucose becomes one of the worst carbon sources for E.coli on poor nitrogen sources due to suboptimal levels of cAMP. Sci Rep 6:24834.

38. Iyer S, Le D, Park BR, & Kim M (2018) Distinct mechanisms coordinate transcription and translation under carbon and nitrogen starvation in Escherichia coli. Nat Microbiol 3(6):741–748.

39. Luo Y, Na Z, & Slavoff SA (2018) P-Bodies: Composition, Properties, and Functions. Biochemistry 57(17):2424–2431.

40. Standart N & Weil D (2018) P-Bodies: Cytosolic Droplets for Coordinated mRNA Storage. Trends Genet 34(8):612–626.

41. van Leeuwen W & Rabouille C (2019) Cellular stress leads to the formation of membraneless stress assemblies in eukaryotic cells. Traffic 20(9):623–638.

42. Datsenko KA & Wanner BL (2000) One-step inactivation of chromosomal genes in Escherichia coli K-12 using PCR products. Proceedings of the National Academy of Sciences of the United States of America 97(12):6640–6645.

43. Stracy M, et al. (2015) Live-cell superresolution microscopy reveals the organization of RNA polymerase in the bacterial nucleoid. Proceedings of the National Academy of Sciences of the United States of America 112(32):E4390–4399.

44. Gibson DG, et al. (2009) Enzymatic assembly of DNA molecules up to several hundred kilobases. Nature methods 6(5):343–345.

45. Baba T, et al. (2006) Construction of Escherichia coli K-12 in-frame, single-gene knockout mutants: the Keio collection. Mol Syst Biol 2:2006 0008.

46. Atlas RM (2010) Handbook of Microbiological Media, Fourth Edition (CRC Press).

47. Endesfelder U, et al. (2013) Multiscale spatial organization of RNA polymerase in Escherichia coli. Biophysical journal 105(1):172–181.

48. Sambrook J, Maniatis T, & Fritsch EF (1989) Molecular cloning : a laboratory manual (Cold Spring Harbor Laboratory Press, Cold Spring Harbor, N.Y.) 2nd Ed p v. (various pagings).

49. Guzman LM, Belin D, Carson MJ, & Beckwith J (1995) Tight regulation, modulation, and high-level expression by vectors containing the arabinose PBAD promoter. Journal of bacteriology 177(14):4121–4130.

